# High Spatial Resolution MALDI Imaging Mass Spectrometry of Fresh-Frozen Bone

**DOI:** 10.1101/2021.10.01.462831

**Authors:** Christopher J. Good, Elizabeth K. Neumann, Casey E. Butrico, James E. Cassat, Richard M. Caprioli, Jeffrey M. Spraggins

## Abstract

Bone and bone marrow are vital to mammalian structure, movement, and immunity. These tissues are also commonly subjected to molecular alterations giving rise to debilitating diseases like rheumatoid arthritis, osteoporosis, osteomyelitis, and cancer. Technologies such as matrix-assisted laser desorption/ionization (MALDI) imaging mass spectrometry (IMS) enable the discovery of spatially resolved chemical information in biological tissue samples to help elucidate the complex molecular processes underlying pathology. Traditionally, preparation of native osseous tissue for MALDI IMS has been difficult due to the mineralized composition and heterogenous morphology of the tissue, and compensation for these challenges with decalcification and fixation protocols can remove or delocalize molecular species. Here, sample preparation methods were advanced to enable multimodal MALDI IMS of undecalcified, fresh-frozen murine femurs allowing the distribution of endogenous lipids to be linked to specific tissue structures and cell types. Adhesive-bound bone sections were mounted onto conductive glass slides with a microscopy-compatible glue and freeze-dried to minimize artificial bone marrow damage. Subliming matrix does not induce further bone marrow cracking, and recrystallizing the deposited matrix improves lipid signal. High spatial resolution (10 μm) MALDI IMS was employed to characterize lipid distributions in fresh-frozen bone, and use of complementary microscopy modalities aided tissue and cell assignments. For example, various phosphatidylcholines localize to bone marrow, adipose tissue, marrow adipose tissue, and muscle. Further, sphingomyelin(42:1) was abundant in megakaryocytes, whereas sphingomyelin(42:2) was diminished in this cell type. These data reflect the vast molecular and cellular heterogeneity indicative of the bone marrow and the soft tissue surrounding the femur. Multimodal MALDI IMS has the potential to advance bone-related biomedical research by offering deep molecular coverage with spatial relevance in a preserved native bone microenvironment.

## INTRODUCTION

Bones provide structural support, enable muscle-mediated mobility, and are responsible for regulating mineral homeostasis, acid-base physiology, and the hematopoietic stem cell niche.^1^ Housed in the intramedullary cavity of long bones, bone marrow possesses extensive cellular and chemical heterogeneity due to hematopoiesis and helps in the development of adaptive and innate immunity. Bone marrow and its skeletal encasing can be compromised by a variety of disorders including those caused by autoinflammation,^2^ bone remodeling imbalances,^3^ nutritional deficiencies,^4^ infection,^5^ and cancer.^6^ Approaches using cell culture, flow cytometry, immunohistochemistry, micro-computed tomography, magnetic resonance imaging have contributed significantly to research on bone-associated diseases.^7–11^ Matrix-assisted laser desorption/ionization (MALDI) imaging mass spectrometry (IMS) is an emerging technology that has the ability to advance our fundamental understanding of bone biology, especially under pathological conditions, by probing the molecular environment in the bone while preserving the native spatial context.

MALDI IMS offers spatially resolved, multi-omics data with high sensitivity and label-free chemical specificity.^12–15^ In brief, laser irradiation is used to ionize endogenous molecules from a tissue surface producing mass spectra in an array of distinct x-y coordinates. The signal intensity of each detected molecule with a given mass-to-charge ratio (*m/*z) can be visualized as a heat map across this surface array. MALDI IMS of bone presents certain challenges resulting from the mineralized and vascularized nature of the tissue. Demineralization strategies have been used to remove the mineral content that can interfere with the detection of organic constituents.^16^ However, these methods can lead to inconsistent preservation of tissue and loss of chemical integrity in terms of the native structure and distribution of biomolecules.^17,18^ Formalin fixation maintains vascular and cellular structures in the bone marrow but results in removal, crosslinking, or delocalization of molecular species. Therefore, imaging of undecalcified and unfixed tissue is ideal, but extensive preparation is required to preserve the heterogenous morphology of the cortical bone, trabecular bone, and bone marrow, all of which differ in porosity and density. Thus, only a handful of studies perform MALDI IMS of fresh-frozen osseous tissue as opposed to fixed and/or decalcified tissue.^19–22^

Seeley et al.^19^ reported on the application of MALDI IMS to a murine cancer model to identify differential expression of proteins within the skeletal environment following tumor growth. In another study, lipids in human bone sections were detected and identified by atmospheric pressure scanning microprobe MALDI IMS.^20^ Further, registration of MALDI generated ion images of lipids with elemental distributions in sections of chicken phalanges has been shown.^21^ In all three studies, an adhesive was utilized to maintain the structural integrity of the undecalcified tissue and assist with mounting mouse, human, or chicken osseous tissue to the MALDI target substrate. For example, Kawomoto et al.^23^ established a method to transfer bone material directly from a tissue block to an adhesive film (Cryofilm) to prevent cryosectioning artifacts, such as cracks and folds in the tissue. Vandenbosch et al.^22^ explored optimal embedding media and matrices for imaging lipids and metabolites in murine femurs. While the results of these studies are encouraging, further investigation of strategies to improve lipid sensitivity and prevent artificial cracking of bone marrow tissue are required if high spatial resolution imaging modalities are to be employed.

Herein, we describe a method to prepare undecalcified, fresh-frozen murine femurs for high spatial resolution (10 μm) MALDI IMS and complementary microscopy. Bone sections bound to an adhesive film are mounted flat to MALDI compatible surfaces by means of a liquid adhesive that possesses minimum interference for autofluorescence and brightfield microscopy and is electrically conductive. Artificial bone marrow damage is assessed for different sample thawing methods and matrix applications, and strategies to increase lipid signal are subsequently investigated. Although many lipid classes are detected in bone-associated tissues and cells, observed phosphatidylcholine (PC) and sphingomyelin (SM) distributions are highlighted. The protocol described herein addresses various challenges that have hindered previous methods for MALDI IMS of fresh-frozen bone tissue and enables high spatial resolution multimodal molecular imaging of this critical organ system.

## METHODS

### Chemicals

Acetone, acetonitrile, hematoxylin, eosin, carboxymethyl cellulose (CMC), and 1,5-diaminonapthalene (DAN, 97%-further purified by recrystallization) were purchased from Sigma-Aldrich (St. Louis, MO, USA). Ethanol, isopentane, isopropanol (IPA), gelatin, and optimal cutting temperature (OCT) compound were purchased from Fisher Scientific (Pittsburgh, PA, USA). Clearium mounting medium was purchased from Electron Microscopy Sciences (Hatfield, PA, USA).

### Tissue Preparation

The right femurs of three 9-week old female C57BL/6J mice (Jackson Laboratory, Bar Harbor, ME, USA) were removed and used for MALDI IMS experiments. Two weeks prior to removal, the mice underwent a procedure on the left leg as part of a separate study; the left femurs were directly inoculated with phosphate buffered saline as detailed in a previously described murine osteomyelitis model.^10^ All animal handling and experimental procedures were conducted in accordance with protocols approved by the Vanderbilt University Institutional Animal Care and Use Committee (IACUC).

Once dissected, femurs were immediately snap frozen over a dry ice-isopentane slurry and stored at −80 °C. Femurs were subsequently embedded in a warm solution of 5% CMC and 10% gelatin^24^ and snap frozen once again over dry ice-isopentane. Embedded femurs were mounted to a chuck using OCT compound. Femurs were cryosectioned at 8 μm thickness using a DB80 HS microtome blade in a CM3050 S cryostat (Leica Biosystems, Wetzlar, Germany) set to −25 °C. Cryofilm 3C 16UF (SECTION-LAB, Hiroshima, Japan) was directly applied to the tissue block to assist in complete tissue transfer.^23^ Cryofilm-bound tissue was mounted flat onto glass slides (Fisher Scientific, Pittsburgh, PA, USA) or conductive indium tin oxide (ITO) coated glass slides (Delta Technologies, Loveland, CO, USA) by means of ZIG 2 way glue (Kuretake Co, Nara, Japan).^20^ For preparations involving thaw-mounting, sections were removed from the cryostat to which heat was directly added, followed by −80 °C storage until being brought to ~22 °C (ambient temperature) in a desiccator. For preparations incorporating freeze-drying, sections were directly stored at −80 °C from the cryostat. Frozen sections were then placed in a glass apparatus surrounded by dry ice and kept under vacuum at ~1 Torr for 8 hours. The chamber was subjected to ambient temperatures for an additional hour. A 7811020 CentriVap Cold Trap (Labconco, Kansas City, MO, USA) was used to condense water vapor.

### Autofluorescence Microscopy

An AxioScan.Z1 slide scanner (Carl Zeiss Microscopy GmbH, Oberkochen, Germany) was used for acquiring autofluorescence images using the EGFP (ex: 488 nm, em: 509 nm), DAPI (ex: 353 nm, em: 465 nm), and DsRed (ex: 545 nm, em: 572 nm) LEDs prior to MALDI IMS. A z-stack of five slices over a 35 μm range was acquired at every tile and merged to account for height differences between the bone marrow and cortical bone. After MALDI IMS acquisition, autofluorescence images were collected using EGFP and brightfield to visualize the MALDI burn patterns and progression of bone marrow cracks.

### Matrix Application

For spraying matrix, a robotic aerosol TM Sprayer (HTX Technologies, Chapel Hill, NC, USA) deposited a 10 mg/mL solution of DAN in 90% acetonitrile onto the sample. Pure acetonitrile flow and N_2_ flow was set to 0.1 mL/min and 10 psi, respectively. The nozzle was heated to 75 °C and allowed to pass across the sample six times with a track spacing of 1.5 mm. The amount of deposited matrix per unit area was measured as 1.6 μg/mm^2^. Alternatively, an in-house sublimation apparatus was used for matrix sublimation.^25,26^ A solution of DAN in acetone (20 mg/mL x 3 mL) was uniformly dispersed on a heated surface and was sublimed at temperatures >130 °C for 10 mins under 30 mTorr vacuum, which resulted in a uniform matrix layer (4.1 μg/mm^2^) on the chilled sample. Following sublimation, the deposited matrix was recrystallized using a hydration chamber as described in a previous method.^27^ In brief, samples were mounted to a metal disc using thermally conductive copper tape (Electron Microscopy Sciences, Hatfield, PA, USA), primed in a 55 °C oven for 2 mins, sealed in a petri dish with 1 mL 5% IPA on filter paper,^28^ heated at 55 °C for 30 s or 90 s, and dried in the 55 °C oven for an additional 2 mins.

### MALDI Q-TOF IMS

A timsTOF fleX (Bruker Daltonics, Bremen, Germany) with a dual ESI-MALDI source was used to acquire all MALDI IMS data.^29^ A SmartBeam 3D 10 kHz frequency tripled Nd:YAG laser (355nm) was tuned to provide optimal signal with the beam scan activated to produce a 16 μm^2^ footprint and 8 μm^2^ footprint for 20 μm and 10 μm spatial resolution imaging, respectively. Local laser power was set to 49% or 71% for 20 μm spatial resolution and decreased to 40% for 10 μm (attenuator offset 0%). Shots remained consistent at 200 per pixel for all experiments. The additional height of the mount and Cryofilm was accounted for by a galvo scanner offset focus adjustment. The MS was operated in qTOF mode with TIMS deactivated. ESI-L Tune Mix (Agilent Technologies, Santa Clara, CA, USA) was directly infused for mass calibration prior to data acquisition. Positive ion mode data was acquired from *m/z* 300-1500. Additional MS parameters are included in the Supporting Information (Table S1).

### MALDI FT-ICR IMS

High mass resolution data was acquired on an FT-ICR MS system from a portion of thaw-mounted bone marrow to obtain high mass accuracy for lipid assignment. A 15T solariX FT-ICR mass spectrometer (Bruker Daltonics, Bremen, Germany) equipped with a Smartbeam II 2 kHz frequency tripled Nd:YAG laser (355 nm) was used. The laser was focused on tissue by adjusting the z-position of the MALDI plate. Laser power, shot count, and pixel size were set to 34%, 275, and 100 μm^2^, respectively. The instrument was mass calibrated using red phosphorus followed by online lock mass calibration to [PC(34:1) + H]^+^ (*m/z* 760.5851). Positive ion mode data was acquired from *m/z* 345-2000 with a time domain file size of 1M, which resulted in a mass resolving power of ~140,000 at *m/z* 760.6. Additional MS parameters are included in the Supporting Information (Table S2).

### Histology

Following post-IMS autofluorescence microscopy, DAN was removed with a pure ethanol wash for 2 mins. The sample was stained with hematoxylin and eosin before a cover slip was mounted with clearium mounting medium. A brightfield microscopy z-stack image was collected at 20x magnification using the Zeiss AxioScan.Z1 slide scanner.

### Data Analysis

All imaging data were visualized using SCiLS Lab Version 2021 (Bruker Daltonics, Bremen, Germany). Ion images were generated using the hot-spot removal function without normalization and denoising. Regions of interest were drawn around the bone marrow using inherent tissue contrast from ion images. These intramedullary cavity regions were used to compare signal between samples. All boxplots were derived from SCiLS by plotting an intensity value for each pixel. Mean intensity values were extracted from peaks in the average mass spectrum of a given region. A peak list was generated and matched with lipids from the LIPIDMAPS database to identify intense ions based on mass accuracy with a <1 ppm mass error tolerance (Table S3).^30,31^

## RESULTS AND DISCUSSION

### Tissue Preparation for Multimodal Molecular Imaging

Cryofilm 3C 16UF maintained its adhesive properties at −25 °C (cryosectioning temperature) and enabled reproducible murine femur sections of 8 μm thickness (Figure 1A). Cryosectioning mineralized and unfixed bone without an adhesive led to tissue loss and distortion that is visible without magnification (Figure S1). For high spatial resolution analysis, it is imperative that the tissue maintains its structure to preserve the spatial accuracy of MALDI IMS data. After adhering the tissue section, the single-sided Cryofilm was mounted flat onto a glass slide using an optically transparent glue that has minimal fluorescence interference at wavelengths used with DAPI, DsRed, and EGFP (Figure 1A). Substrate and adhesive transparency are required for multimodal imaging experiments where brightfield and fluorescence microscopy can be used to annotate the functional structures and cell types within the tissue. If the mount inherently produces fluorescence background, it would be more challenging to identify morphological structures without additional stains, many of which are incompatible with mass spectrometry analysis. Furthermore, a mount that minimizes air bubbles and promotes a level tissue surface is necessary for MALDI IMS to limit topographical changes that can lead to defocusing of the laser at the tissue surface. Overall, the Cryofilm and mounting adhesive are ideal for generating reproducible bone samples that are compatible with MALDI IMS and microscopy modalities.

**Figure 1.**
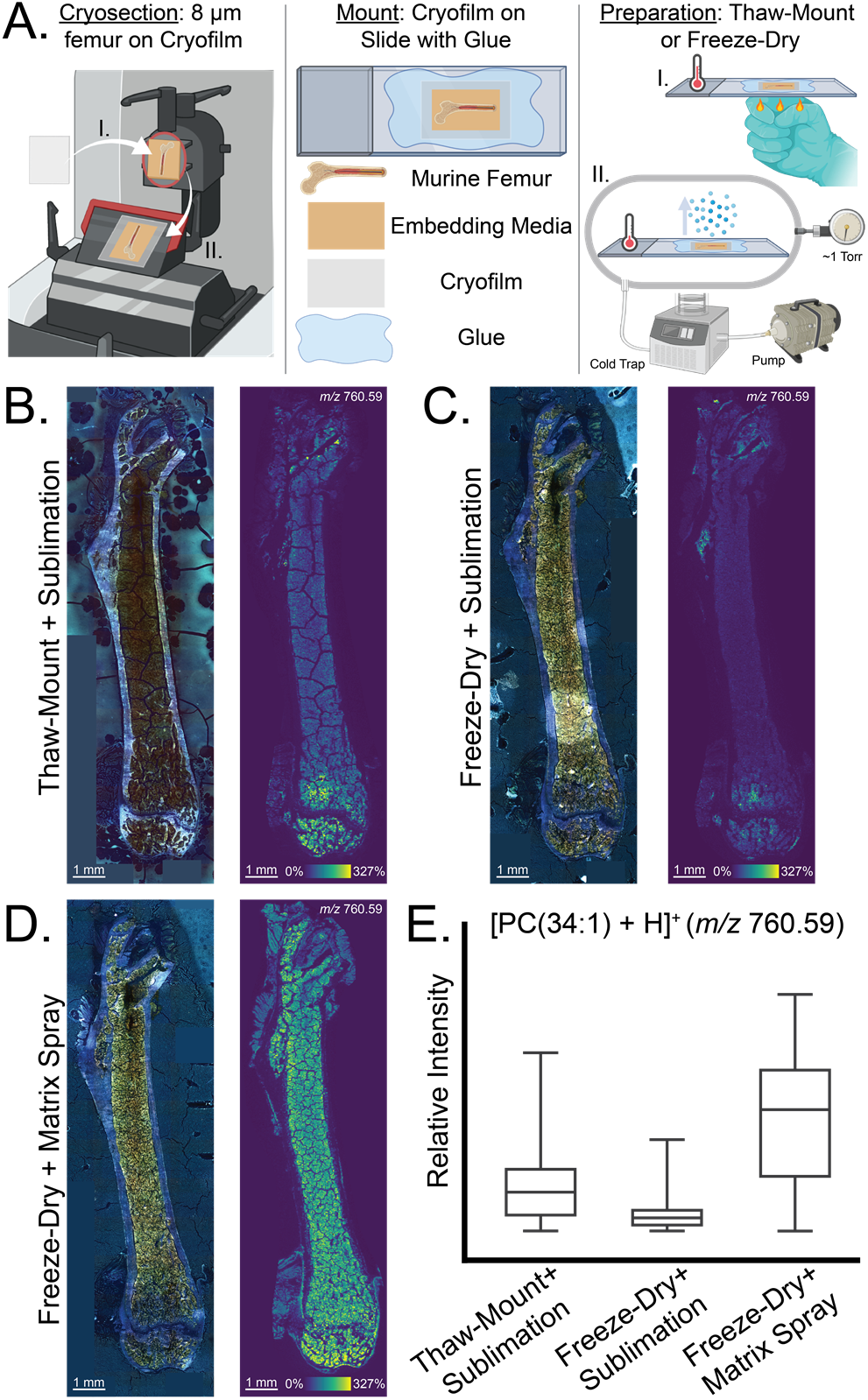
Methods for preparing adhesive-bound, fresh-frozen femur sections lead to differences in bone marrow morphology and signal intensity of biomolecules. (A) Overview of sample preparation that is compatible with MALDI IMS and microscopy modalities: cryosectioning (I. apply film, II. cut section), mounting, and preparing sections (I. thaw-mount, II. freeze-dry). (B, C, D) Merged channel autofluorescence images (left) and ion images of [PC(34:1) + H]^+^ (*m/z* 760.59; right) highlight morphological damage and lipid signal between thaw-mounting, freeze-drying, matrix sublimation, and matrix spraying. (B) Thaw-mounting and subliming matrix yield large bone marrow cracks. (C) Freeze-drying and subliming matrix result in less pronounced cracks but worse signal. (D) Freeze-drying and spraying matrix recover signal with an additional sacrifice to morphology. (E) Boxplots recapitulate intensity differences of [PC(34:1) + H]^+^ (*m/z* 760.59) between sample preparation methods. Boxplots represent the range and interquartile range of a molecule’s intensity values, which are derived from all pixels within a singular intramedullary cavity. These MALDI IMS data were collected in positive ion mode at 20 μm spatial resolution. Ion images were not normalized for intensity comparisons between tissue sections.

### Balancing Tissue Morphology and Signal Intensity When Thawing and Applying Matrix

Strategies to bring frozen femur sections to ambient temperatures (~22 °C) were investigated (Figure 1B). A tradeoff arose between preserving bone marrow morphology and achieving adequate signal intensity of biomolecules. Tissue sections are thaw-mounted directly to the slide in traditional MALDI IMS experiments; however, thaw-mounting a femur section directly onto Cryofilm resulted in artificial bone marrow cracks as large as 75 μm wide (Figure S2). We hypothesize that the residual water content in this gelatinous soft tissue is responsible for artifact formation. Removing water content by lyophilization/freeze-drying yielded less pronounced cracks that are nearly indistinguishable by MALDI IMS at 20 μm spatial resolution (Figure 1C), although, cracks were still noticeable when using microscopy modalities. Previously, Saigusa et al.^32^ reported an increase in metabolite signal in rat brain that was thaw-mounted to Cryofilm compared to freeze-dried sections, and we observed the same trend for lipid signal in murine bone marrow (Figure 1B, 1C, 1E). After mounting tissue onto a slide, matrix was deposited using either a robotic aerosol sprayer or sublimation apparatus, and spraying resulted in an intensity increase for lipids such as [PC(34:1) + H]^+^ (*m/z* 760.59, Figure 1D, 1E). The solvents used when depositing matrix aid in the extraction of molecules from the tissue surface, which results in enhanced signal. This is a phenomenon that does not occur with dry matrix applications like sublimation.

Following closer examination of tissue morphology, solvent exposure from spraying matrix contributed to additional bone marrow damage or cracks (Figure 2). Minor artifacts manifested from the freeze-drying process, likely due to incomplete water removal from the tissue. These cracks appear in the pre-IMS images and are accentuated in the post-IMS images, regardless of the matrix application method (Figure 2A, 2B). In the post-IMS autofluorescence images, the matrix layer adds contrast to the artifacts and increases their visibility. Moreover, during H&E staining protocols, ethanol washing shrunk the bone marrow and exacerbated the cracks in the tissue. Brightfield microscopy also reveals bone marrow artifacts because the light source illuminates the sample through the backside of the slide. Nevertheless, tissue exposure to sublimed DAN and MALDI laser irradiation did not significantly alter tissue morphology (Figure 2A). This observation is confirmed in Figure 3C when comparing both pre- and post-IMS brightfield images and is consistent with previous literature.^33^ In contrast, sprayed DAN contributed to additional and more pronounced bone marrow cracks that compromised ion and histological images (Figure 2B). Aerosolized solvent exposure was the source of additional morphological damage, and thus sublimation was determined to be the optimal matrix application method.

**Figure 2.**
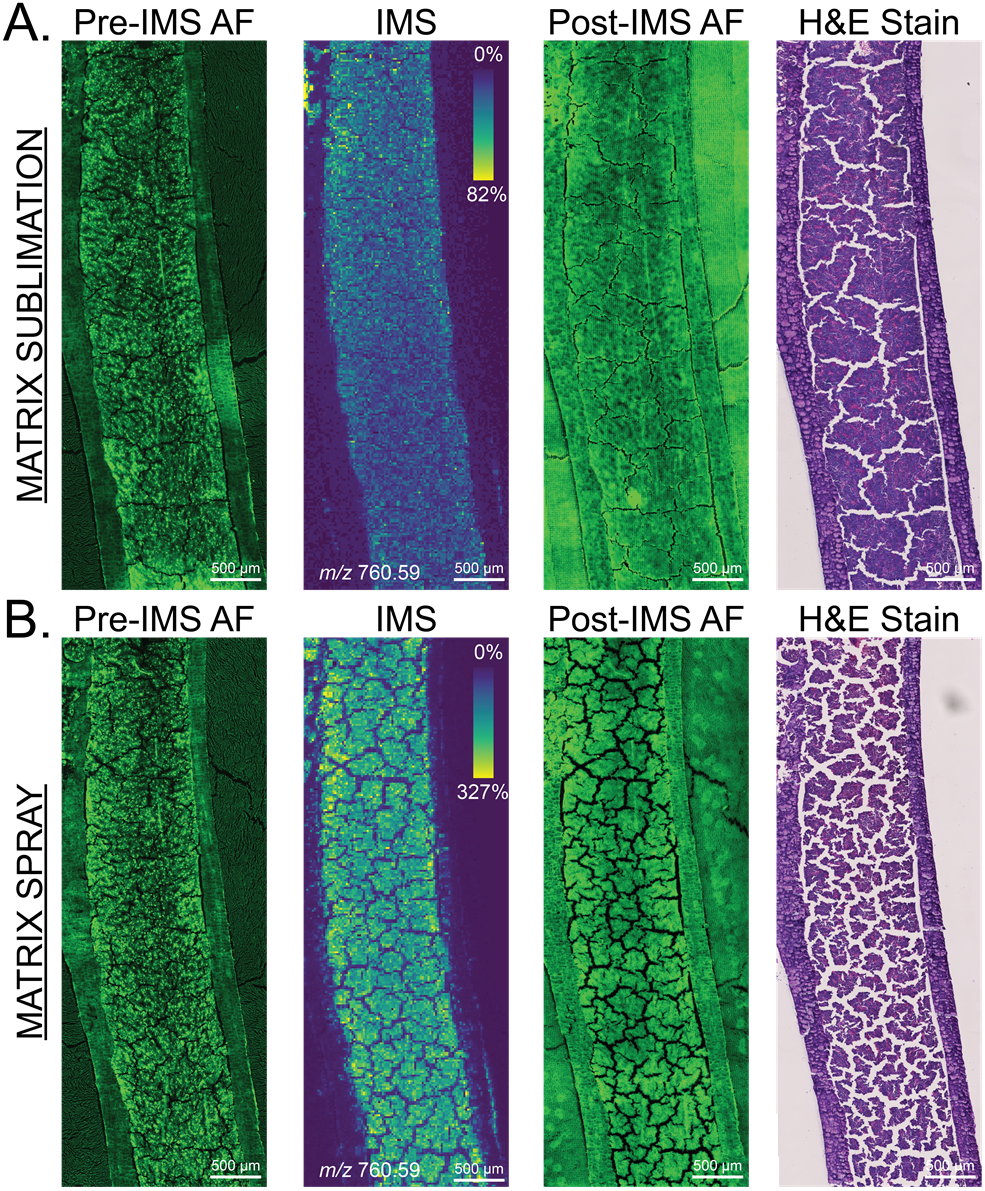
Spraying matrix generates more severe bone marrow artifacts that negatively affect MALDI IMS data and downstream microscopy. (A, B) Autofluorescence (EGFP, pre- and post-IMS), 20 μm spatial resolution ion images of [PC(34:1) + H]^+^ (*m/z* 760.59), and H&E staining display the propagation of bone marrow cracks throughout a MALDI IMS experiment. Bone marrow cracks are less pronounced when (A) matrix is sublimed compared to when (B) matrix is sprayed. The matrix layer appears in the post-IMS autofluorescence images. Ion images were not normalized.

**Figure 3.**
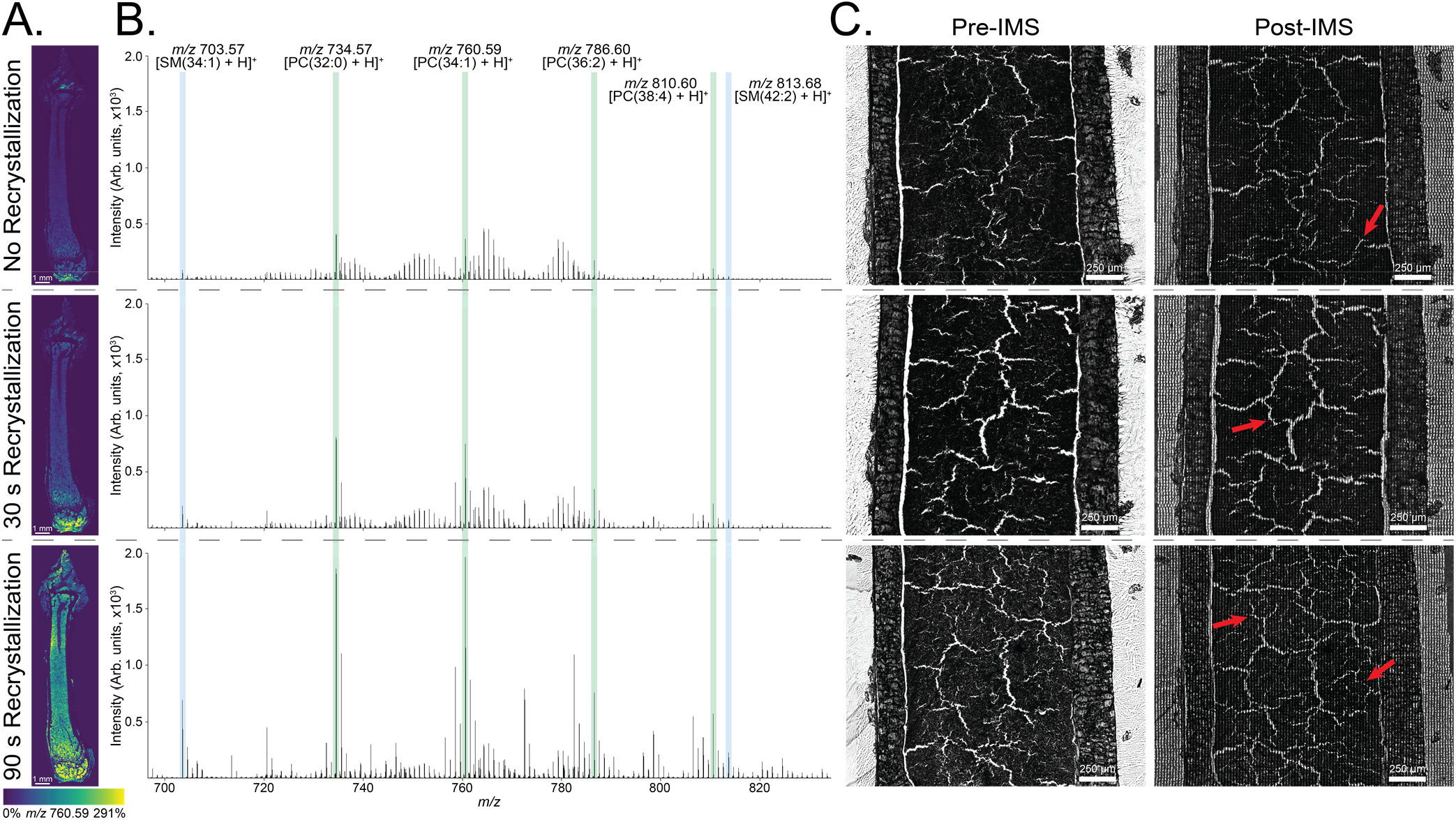
Recrystallizing the deposited matrix increases lipid signal and does not significantly contribute to bone marrow damage. (A, B, C) Images and spectra are aligned across; top represents no recrystallization, middle represents 30 s recrystallization, bottom represents 90 s recrystallization. (A) 20 μm spatial resolution ion images of [PC(34:1) + H]^+^ (*m/z* 760.59) highlight the improvement in signal for full femur sections as recrystallization time increases. (B) Unnormalized mean spectra show an intensity increase for various PCs (green) and SMs (blue). (C) Red arrows in the post-IMS brightfield images indicate minor cracks that manifest from matrix deposition, recrystallization (if applicable), and MALDI IMS.

### Improving Lipid Signal with Matrix Recrystallization and Conductive Coatings

Freeze-drying femur sections and applying matrix by sublimation resulted in the least amount of bone marrow damage; however, the overall lipid signal was relatively low. Recrystallizing the matrix layer with 5% aqueous IPA improved the signal intensity of most lipid species in a time-dependent manner, including [PC(34:1) + H]^+^ (*m/z* 760.59; Figure 3A, 3B). Improvement in signal was observed with as little as 30 s of recrystallization, and 90 s of exposure to IPA vapor yielded a 437% intensity increase for [PC(34:1) + H]^+^ (*m/z* 760.59) with minimal molecular delocalization. This trend was consistent regardless of lipid class, degree of unsaturation, or acyl chain length as observed with [PC(32:0) + H]^+^ (*m/*z 734.57), [PC(36:2) + H]^+^ (*m/*z 786.60), [PC(38:4) + H]^+^ (*m/*z 810.60), [PC(34:1) + H]^+^ (*m/*z 760.59), [SM(34:1) + H]^+^ (*m/*z 703.57), and [SM(42:2) + H]^+^ (*m/z* 813.68; Figure 3B). Additional bone marrow damage did not appear to be significant following the recrystallization procedure (Figure 3C). There was evidence of the formation of <10 μm cracks; however, the number and severity of newly formed cracks did not correlate with different recrystallization time points. We anticipate that a further increase in recrystallization time will improve signal but at the cost of spatial resolution due to molecular diffusion and morphological damage. Optimization of these time points may be required when analyzing different organ systems. Overall, recrystallizing DAN on freeze-dried femur sections enables ample lipid signal without further compromising bone marrow morphology.

Cryofilm 3C 16UF was predicted to be nonconductive since it is comprised of a polyvinylidene chloride film and acrylic resin cryoglue.^34^ For MALDI IMS, a charged surface enhances ion acceleration and dissipates charge accumulation on the tissue surface, which improves ionization and signal. Therefore, alternative conductive adhesives have been explored for imaging mass spectrometry.^22,32^ Interestingly, a notable intensity increase was discovered when the Cryofilm-bound tissue was mounted onto an ITO coated glass slide in lieu of a glass slide (Figure 4). All *m/z* features detected in the intramedullary cavity, including those identified as PCs and SMs, exemplified greater signal when mounted to a conductive surface. This trend held true for lower abundant lipids like [SM(42:2) + H]^+^ (*m/*z 813.68), where a 34.9% intensity increase was still observed. The combination of Cryofilm and mounting adhesive is sufficiently conductive to permit the advantage gained from an ITO coating.

**Figure 4.**
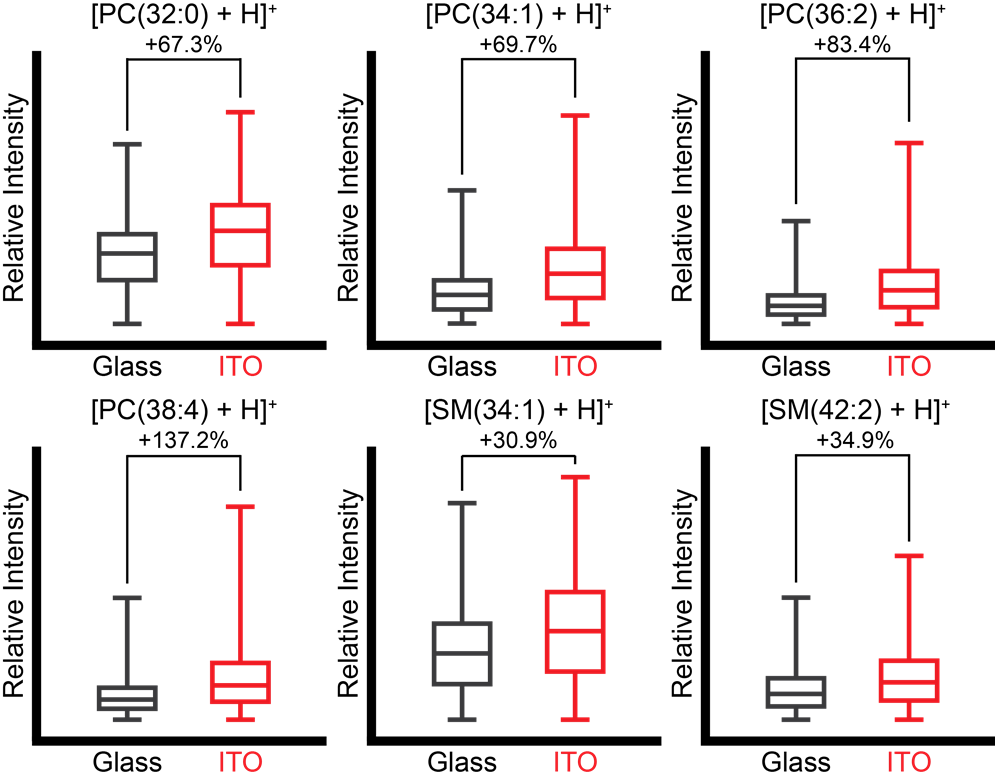
Cryofilm-bound tissue mounted to an ITO coated glass slide provides an additional improvement in signal for all PCs and SMs regardless of their native abundances in tissue. Boxplots represent the range and interquartile range of a molecule’s intensity values, which are derived from all pixels within a singular intramedullary cavity. Please note that the intensity scaling is different for each boxplot, and the intensity increase percentages were calculated from the unnormalized mean intensity values for each region. Deposited matrix was recrystallized for 30 s, and MALDI IMS data were collected at 20 μm spatial resolution.

### Molecular Mapping of Tissue Structures and Immune Cells in Bone Tissue

The sample preparation methods discussed thus far were leveraged to visualize and define endogenous lipid distributions in macroscopic soft tissues within and surrounding the intramedullary cavity of undecalcified bone (Figure 5). Pre-IMS autofluorescence and post-IMS histological staining were used to define functional tissue units, such as cortical bone, trabecular bone, bone marrow, adipose tissue, and muscle. Lipids localized to most soft tissue structures, but some showed differential distributions based on acyl chain length and degree of unsaturation. For example, [PC(32:0) + H]^+^ (*m/*z 734.57) was the most abundant lipid in the bone marrow, while [PC(36:2) + H]^+^ (*m/*z 786.60) and [PC(38:6) + H]^+^ (*m/*z 806.57) primarily distributed in contiguous adipose tissue and muscle tissue, respectively. There was no identifiable lipid signal from the cortical or trabecular bone regions, which is consistent with previously published studies using DAN.^22^

**Figure 5.**
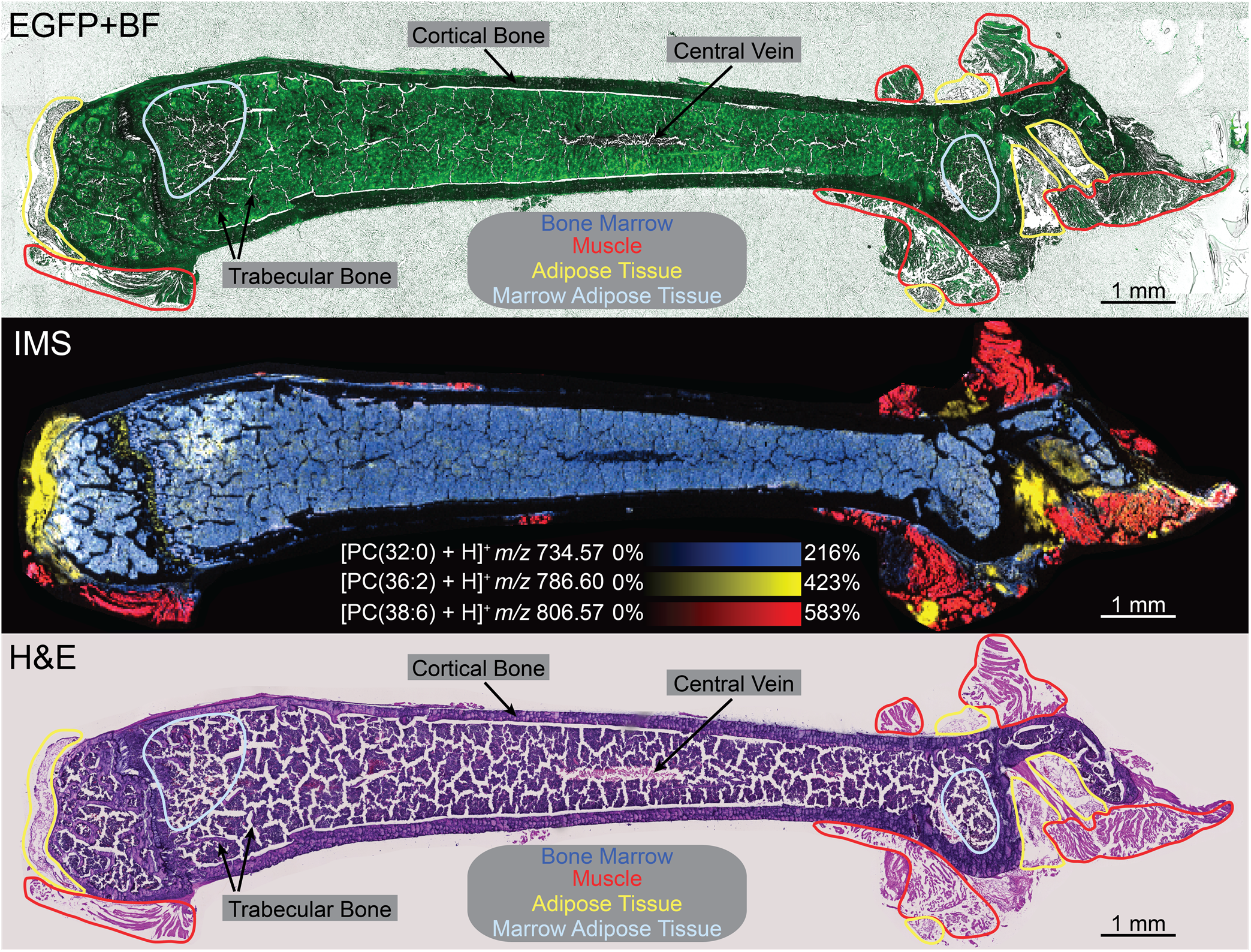
Sample preparation methods enable visualization of lipid distributions in various tissue structures defined by histology. [PC(32:0) + H]^+^ (*m/z* 734.57), [PC(36:2) + H]^+^ (*m/z* 786.60), and [PC(38:6) + H]^+^ (*m/z* 806.57) localize to bone marrow, adipose tissue, and muscle, respectively. Pre-IMS autofluorescence (EGFP) with brightfield (BF) microscopy and post-IMS histological staining (H&E) aid in tissue and cell assignments. MALDI IMS data were collected at 20 μm spatial resolution, and ion images were not normalized.

Adipocytes comprise a central mass of stored triglycerides surrounded by a phospholipid membrane. In addition to surrounding the femur as white adipose tissue, adipocytes constitute a portion of bone marrow cellular content called marrow adipose tissue.^35^ Colocalization of [PC(32:0) + H]^+^ (*m/*z 734.57) and [PC(36:2) + H]^+^ (*m/*z 786.60) was observed at the distal end of the femur. These lipid signatures provide a molecular and visual representation of marrow adipose tissue, which is consistent with previous findings. Marrow adipose tissue volume in long bones of 4-12 week old C57BL/6J mice has been determined to be greater in distal regions compared to proximal regions.^36^ Others have also identified PC(36:2) in white and brown murine adipose tissues,^37^ both of which possess analogous characteristics of marrow adipose tissue.^38^

Our new sample preparation and MALDI IMS technologies enabled imaging of femur sections at 10 μm spatial resolution (Figure S3). Even though bone marrow artifacts are accentuated in the higher spatial resolution images, lipid profiles from individual cells and vasculature were able to be detected with this method (Figure 6). Microscopy highlighted the location of a blood vessel and hematopoietic cells in the bone marrow, and histological analysis suggested the identity of these large cells as megakaryocytes (Figure 6A). Megakaryocytes generate platelets and have diverse cytoplasmic and membrane lipid content that consist of both phosphatidylcholines and sphingomyelins.^39^ As differentiated megakaryocytes mature, their cell diameter expands and the nucleus becomes multilobed.^40^ We have identified megakaryocytes with different diameters (~20-30 μm) and nuclear configurations that exemplify variations in maturity (Figure S4). Regardless of maturity, megakaryocytes were observed to have greater abundances of some sphingomyelins relative to surrounding cell types (Figure 6B). For instance, [SM(42:1) + H]^+^ (*m/z* 815.70) and [SM(40:1) + H]^+^ (*m/z* 787.67) possessed higher abundances within the megakaryocytes. In contrast, [SM(42:2) + H]^+^ (*m/z* 813.68) differs by only one degree of unsaturation and exhibited a decrease in abundance within these cells. High spatial resolution imaging also uncovered specific lipids that localized to blood vessels within the bone, such as [SM(40:1) + H]^+^ (*m/z* 787.67) and [SM(36:1) + H]^+^ (*m/z* 731.61). These data suggest some degree of lipid preservation following the complex molecular process of thrombopoiesis within the bone marrow.

**Figure 6.**
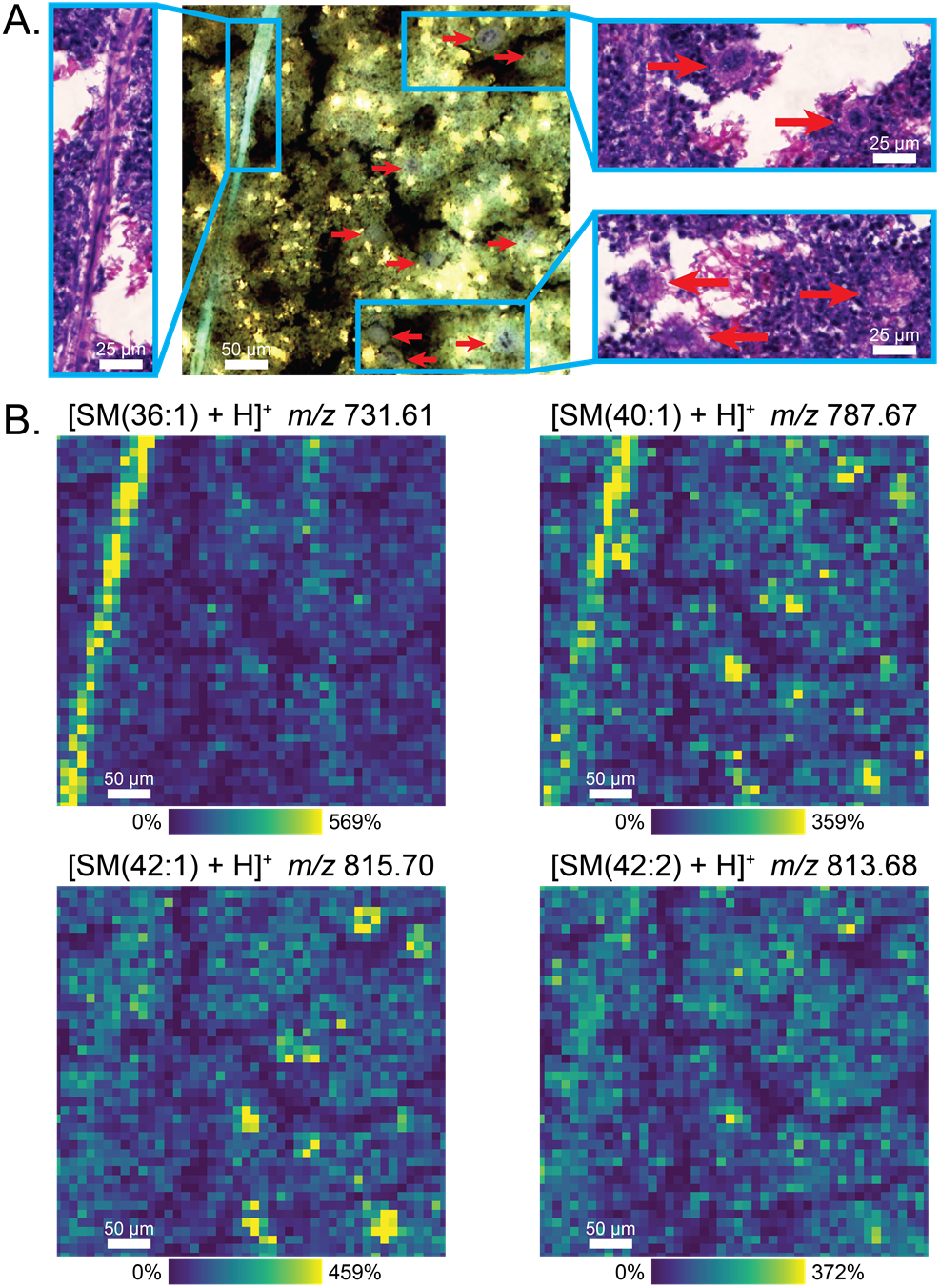
Sphingomyelins localize to microscopic vascular structures and individual megakaryocytes in the bone marrow. (A) Merged channel autofluorescence and enlarged H&E stain images reveal a blood vessel and multiple megakaryocytes (red arrows). (B) [SM(36:1) + H]^+^ (*m/z* 731.61) and [SM(40:1) + H]^+^ (*m/z* 787.67) localize to the blood vessel. The latter also localizes to individual megakaryocytes in addition to [SM(42:1) + H]^+^ (*m/z* 815.70). An extra degree of unsaturation decreases the presence of the 42-carbon chain SM within the cells. MALDI IMS data were collected at 10 μm spatial resolution, and ion images were not normalized.

## CONCLUSIONS

The advanced sample preparation methods described herein allowed us to perform multimodal molecular imaging of undecalcified, fresh-frozen murine femurs at cellular spatial resolution. To achieve optimal signal intensity of biomolecules while preserving unfixed bone marrow morphology, Cryofilm-assisted cryosections were mounted flat onto ITO coated glass slides via an adhesive and subsequently freeze-dried. The matrix sublimation and recrystallization processes do not exacerbate bone marrow damage and yield ample lipid signal. We were able to identify lipids that localize to bone marrow, adipose tissue, marrow adipose tissue, muscle, blood vessels, and megakaryocytes. We also determined that chain length and degree of unsaturation of sphingomyelins dictate their abundances within megakaryocytes. Tissue and cell assignments were made using complementary microscopy modalities, which emphasizes the importance of developing methods that are compatible with these imaging technologies. We believe this multimodal MALDI IMS workflow can be leveraged for research of metabolic processes associated with marrow adipose tissue, physiological mechanisms like thrombopoiesis, and molecular modifications resulting from pathological conditions within the bone microenvironment.

## ACKNOWLEDGEMENTS

The authors would like to thank Dr. Eric Spivey and Dr. David Anderson for developing and troubleshooting the in-house sublimation apparatus. Figure 1A was created with BioRender.com (BQ231UWMH9). Support was provided by the NIH National Institute of Allergy and Infectious Disease (R01 AI145992 awarded to J.M.S. and J.E.C.), NIH Shared Instrumentation Grant Program (S10 OD012359 awarded to R.M.C.), the National Science Foundation Major Research Instrument Program (CBET 1828299 awarded to J.M.S. and R.M.C.). C.J.G. and C.E.B. are supported by the NIH National Institute of Allergy and Infectious Disease (T32 AI112541). E.K.N. is supported by the National Institute of Environmental Health Sciences training grant (T32 ES007028). J.E.C. is also supported by the NIH National Institute of Allergy and Infectious Disease (R01 AI132560) and a Burroughs Welcome Fund Career Award for Medical Scientists.

---

**TOC FIGURE.**
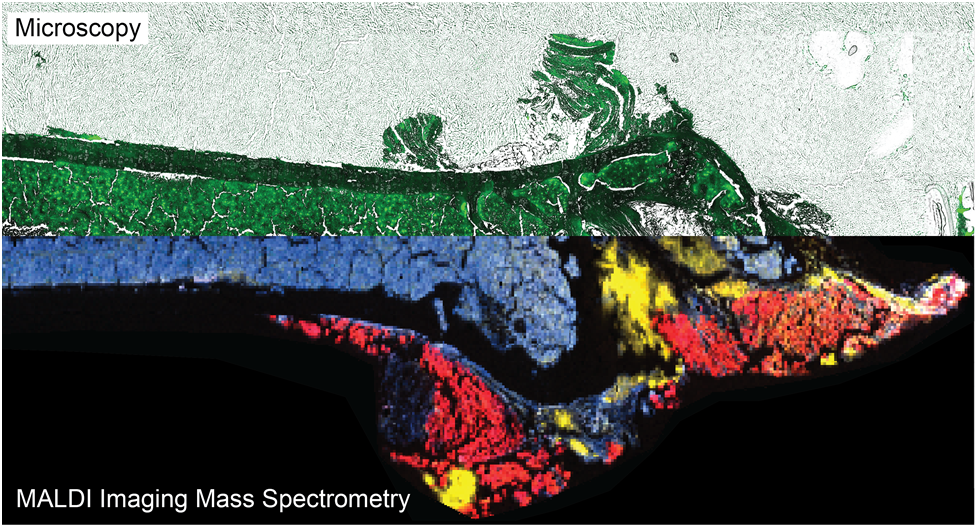

## REFERENCES

(1) Clarke, B. Normal Bone Anatomy and Physiology. Clin J Am Soc Nephrol 2008, 3 (Suppl 3), S131–S139. https://doi.org/10.2215/CJN.04151206.

(2) Ferguson, P. J.; El-Shanti, H. I. Autoinflammatory Bone Disorders. Current Opinion in Rheumatology 2007, 19 (5), 492–498. https://doi.org/10.1097/BOR.0b013e32825f5492.

(3) Feng, X.; McDonald, J. M. Disorders of Bone Remodeling. Annual Review of Pathology: Mechanisms of Disease 2011, 6 (1), 121–145. https://doi.org/10.1146/annurev-pathol-011110-130203.

(4) Karpouzos, A.; Diamantis, E.; Farmaki, P.; Savvanis, S.; Troupis, T. Nutritional Aspects of Bone Health and Fracture Healing. J Osteoporos 2017, 2017, 4218472. https://doi.org/10.1155/2017/4218472.

(5) Lew, D. P.; Waldvogel, F. A. Osteomyelitis. The Lancet 2004, 364 (9431), 369–379. https://doi.org/10.1016/S0140-6736(04)16727-5.

(6) Rajkumar, S. V.; Kumar, S. Multiple Myeloma: Diagnosis and Treatment. Mayo Clinic Proceedings 2016, 91 (1), 101–119. https://doi.org/10.1016/j.mayocp.2015.11.007.

(7) Kuca-Warnawin, E.; Kurowska, W.; Prochorec-Sobieszek, M.; Radzikowska, A.; Burakowski, T.; Skalska, U.; Massalska, M.; Plebańczyk, M.; Małdyk-Nowakowska, B.; Słowińska, I.; Gasik, R.; Maśliński, W. Rheumatoid Arthritis Bone Marrow Environment Supports Th17 Response. Arthritis Res Ther 2017, 19, 274. https://doi.org/10.1186/s13075-017-1483-x.

(8) Aalbers, A. M.; van den Heuvel-Eibrink, M. M.; Baumann, I.; Dworzak, M.; Hasle, H.; Locatelli, F.; Moerloose, B. D.; Schmugge, M.; Mejstrikova, E.; Nováková, M.; Zecca, M.; Zwaan, C. M.; te Marvelde, J. G.; Langerak, A. W.; van Dongen, J. J. M.; Pieters, R.; Niemeyer, C. M.; van der Velden, V. H. J.. Bone Marrow Immunophenotyping by Flow Cytometry in Refractory Cytopenia of Childhood. Haematologica 2015, 100 (3), 315–323. https://doi.org/10.3324/haematol.2014.107706.

(9) Bussard, K. M.; Okita, N.; Sharkey, N.; Neuberger, T.; Webb, A.; Mastro, A. M. Localization of Osteoblast Inflammatory Cytokines MCP-1 and VEGF to the Matrix of the Trabecula of the Femur, a Target Area for Metastatic Breast Cancer Cell Colonization. Clin Exp Metastasis 2010, 27 (5), 331–340. https://doi.org/10.1007/s10585-010-9330-3.

(10) Cassat, J. E.; Hammer, N. D.; Campbell, J. P.; Benson, M. A.; Perrien, D. S.; Mrak, L. N.; Smeltzer, M. S.; Torres, V. J.; Skaar, E. P. A Secreted Bacterial Protease Tailors the Staphylococcus Aureus Virulence Repertoire to Modulate Bone Remodeling during Osteomyelitis. Cell Host Microbe 2013, 13 (6), 759–772. https://doi.org/10.1016/j.chom.2013.05.003.

(11) Daffner, R.; Lupetin, A.; Dash, N.; Deeb, Z.; Sefczek, R.; Schapiro, R. MRI in the Detection of Malignant Infiltration of Bone Marrow. American Journal of Roentgenology 1986, 146 (2), 353–358. https://doi.org/10.2214/ajr.146.2.353.

(12) Caprioli, R. M.; Farmer, T. B.; Gile, J. Molecular Imaging of Biological Samples: Localization of Peptides and Proteins Using MALDI-TOF MS. Anal. Chem. 1997, 69 (23), 4751–4760. https://doi.org/10.1021/ac970888i.

(13) Stoeckli, M.; Chaurand, P.; Hallahan, D. E.; Caprioli, R. M. Imaging Mass Spectrometry: A New Technology for the Analysis of Protein Expression in Mammalian Tissues. Nature Medicine 2001, 7 (4), 493–496. https://doi.org/10.1038/86573.

(14) McDonnell, L. A.; Heeren, R. M. A. Imaging Mass Spectrometry. Mass Spectrometry Reviews 2007, 26 (4), 606–643. https://doi.org/10.1002/mas.20124.

(15) Karlsson, O.; Hanrieder, J. Imaging Mass Spectrometry in Drug Development and Toxicology. Arch. Toxicol. 2017, 91 (6), 2283–2294. https://doi.org/10.1007/s00204-016-1905-6.

(16) Hirano, H.; Masaki, N.; Hayasaka, T.; Watanabe, Y.; Masumoto, K.; Nagata, T.; Katou, F.; Setou, M. Matrix-Assisted Laser Desorption/Ionization Imaging Mass Spectrometry Revealed Traces of Dental Problem Associated with Dental Structure. Anal Bioanal Chem 2014, 406 (5), 1355–1363. https://doi.org/10.1007/s00216-013-7075-y.

(17) Hosoya, A.; Hoshi, K.; Sahara, N.; Ninomiya, T.; Akahane, S.; Kawamoto, T.; Ozawa, H. Effects of Fixation and Decalcification on the Immunohistochemical Localization of Bone Matrix Proteins in Fresh-Frozen Bone Sections. Histochem Cell Biol 2005, 123 (6), 639–646. https://doi.org/10.1007/s00418-005-0791-4.

(18) Liu, H.; Zhu, R.; Liu, C.; Ma, R.; Wang, L.; Chen, B.; Li, L.; Niu, J.; Zhao, D.; Mo, F.; Fu, M.; Brömme, D.; Zhang, D.; Gao, S. Evaluation of Decalcification Techniques for Rat Femurs Using HE and Immunohistochemical Staining. BioMed Research International 2017, 2017, e9050754. https://doi.org/10.1155/2017/9050754.

(19) Seeley, E. H.; Wilson, K. J.; Yankeelov, T. E.; Johnson, R. W.; Gore, J. C.; Caprioli, R. M.; Matrisian, L. M.; Sterling, J. A. Co-Registration of Multi-Modality Imaging Allows for Comprehensive Analysis of Tumor-Induced Bone Disease. Bone 2014, 61, 208–216. https://doi.org/10.1016/j.bone.2014.01.017.

(20) Schaepe, K.; Bhandari, D. R.; Werner, J.; Henss, A.; Pirkl, A.; Kleine-Boymann, M.; Rohnke, M.; Wenisch, S.; Neumann, E.; Janek, J.; Spengler, B. Imaging of Lipids in Native Human Bone Sections Using TOF– Secondary Ion Mass Spectrometry, Atmospheric Pressure Scanning Microprobe Matrix-Assisted Laser Desorption/Ionization Orbitrap Mass Spectrometry, and Orbitrap–Secondary Ion Mass Spectrometry. 2018. https://doi.org/10.1021/acs.analchem.8b00892.s001.

(21) Svirkova, A.; Turyanskaya, A.; Perneczky, L.; Streli, C.; Marchetti-Deschmann, M. Multimodal Imaging of Undecalcified Tissue Sections by MALDI MS and ΜXRF. Analyst 2018, 143 (11), 2587–2595. https://doi.org/10.1039/c8an00313k.

(22) Vandenbosch, M.; Nauta, S. P.; Svirkova, A.; Poeze, M.; Heeren, R. M. A.; Siegel, T. P.; Cuypers, E.; Marchetti-Deschmann, M. Sample Preparation of Bone Tissue for MALDI-MSI for Forensic and (Pre)Clinical Applications. Anal Bioanal Chem 2020. https://doi.org/10.1007/s00216-020-02920-1.

(23) Kawamoto, T.; Kawamoto, K. Preparation of Thin Frozen Sections from Nonfixed and Undecalcified Hard Tissues Using Kawamot’s Film Method (2012). In Skeletal Development and Repair: Methods and Protocols; Hilton, M. J., Ed.; Methods in Molecular Biology; Humana Press: Totowa, NJ, 2014; pp 149–164. https://doi.org/10.1007/978-1-62703-989-5_11.

(24) Nelson, K. A.; Daniels, G. J.; Fournie, J. W.; Hemmer, M. J. Optimization of Whole-Body Zebrafish Sectioning Methods for Mass Spectrometry Imaging. J Biomol Tech 2013, 24 (3), 119–127. https://doi.org/10.7171/jbt.13-2403-002.

(25) Hankin, J. A.; Barkley, R. M.; Murphy, R. C. Sublimation as a Method of Matrix Application for Mass Spectrometric Imaging. J Am Soc Mass Spectrom 2007, 18 (9), 1646–1652. https://doi.org/10.1016/j.jasms.2007.06.010.

(26) Thomas, A.; Charbonneau, J. L.; Fournaise, E.; Chaurand, P. Sublimation of New Matrix Candidates for High Spatial Resolution Imaging Mass Spectrometry of Lipids: Enhanced Information in Both Positive and Negative Polarities after 1,5-Diaminonapthalene Deposition. Anal. Chem. 2012, 84 (4), 2048–2054. https://doi.org/10.1021/ac2033547.

(27) Yang, J.; Caprioli, R. M. Matrix Sublimation/Recrystallization for Imaging Proteins by Mass Spectrometry at High Spatial Resolution. Anal. Chem. 2011, 83 (14), 5728–5734. https://doi.org/10.1021/ac200998a.

(28) Dueñas, M. E.; Carlucci, L.; Lee, Y. J. Matrix Recrystallization for MALDI-MS Imaging of Maize Lipids at High-Spatial Resolution. J. Am. Soc. Mass Spectrom. 2016, 27 (9), 1575–1578. https://doi.org/10.1007/s13361-016-1422-0.

(29) Spraggins, J. M.; Djambazova, K. V.; Rivera, E. S.; Migas, L. G.; Neumann, E. K.; Fuetterer, A.; Suetering, J.; Goedecke, N.; Ly, A.; Van de Plas, R.; Caprioli, R. M. High-Performance Molecular Imaging with MALDI Trapped Ion-Mobility Time-of-Flight (TimsTOF) Mass Spectrometry. Anal. Chem. 2019, 91 (22), 14552–14560. https://doi.org/10.1021/acs.analchem.9b03612.

(30) Sud, M.; Fahy, E.; Cotter, D.; Brown, A.; Dennis, E. A.; Glass, C. K.; Merrill, A. H.; Murphy, R. C.; Raetz, C. R. H.; Russell, D. W.; Subramaniam, S. LMSD: LIPID MAPS Structure Database. Nucleic Acids Res 2007, 35 (Database issue), D527–D532. https://doi.org/10.1093/nar/gkl838.

(31) Liebisch, G.; Fahy, E.; Aoki, J.; Dennis, E. A.; Durand, T.; Ejsing, C. S.; Fedorova, M.; Feussner, I.; Griffiths, W. J.; Köfeler, H.; Merrill, A. H.; Murphy, R. C.; O’Donnell, V. B.; Oskolkova, O.; Subramaniam, S.; Wakelam, M. J. O.; Spener, F. Update on LIPID MAPS Classification, Nomenclature, and Shorthand Notation for MS-Derived Lipid Structures. Journal of Lipid Research 2020, 61 (12), 1539–1555. https://doi.org/10.1194/jlr.S120001025.

(32) Saigusa, D.; Saito, R.; Kawamoto, K.; Uruno, A.; Kano, K.; Aoki, J.; Yamamoto, M.; Kawamoto, T. Conductive Adhesive Film Expands the Utility of Matrix-Assisted Laser Desorption/Ionization Mass Spectrometry Imaging. Anal. Chem. 2019, 91 (14), 8979–8986. https://doi.org/10.1021/acs.analchem.9b01159.

(33) Neumann, E. K.; Comi, T. J.; Rubakhin, S. S.; Sweedler, J. V. Lipid Heterogeneity between Astrocytes and Neurons Revealed by Single-Cell MALDI-MS Combined with Immunocytochemical Classification. Angewandte Chemie 2019, 131 (18), 5971–5975. https://doi.org/10.1002/ange.201812892.

(34) Ticha, P.; Pilawski, I.; Yuan, X.; Pan, J.; Tulu, U. S.; Coyac, B. R.; Hoffmann, W.; Helms, J. A. A Novel Cryo-Embedding Method for in-Depth Analysis of Craniofacial Mini Pig Bone Specimens. Sci Rep 2020, 10 (1), 19510. https://doi.org/10.1038/s41598-020-76336-3.

(35) Li, Y.; Meng, Y.; Yu, X. The Unique Metabolic Characteristics of Bone Marrow Adipose Tissue. Frontiers in Endocrinology 2019, 10, 69. https://doi.org/10.3389/fendo.2019.00069.

(36) Scheller, E. L.; Doucette, C. R.; Learman, B. S.; Cawthorn, W. P.; Khandaker, S.; Schell, B.; Wu, B.; Ding, S.-Y.; Bredella, M. A.; Fazeli, P. K.; Khoury, B.; Jepsen, K. J.; Pilch, P. F.; Klibanski, A.; Rosen, C. J.; MacDougald, O. A. Region-Specific Variation in the Properties of Skeletal Adipocytes Reveals Regulated and Constitutive Marrow Adipose Tissues. Nat Commun 2015, 6 (1), 7808. https://doi.org/10.1038/ncomms8808.

(37) Hoene, M.; Li, J.; Häring, H.-U.; Weigert, C.; Xu, G.; Lehmann, R. The Lipid Profile of Brown Adipose Tissue Is Sex-Specific in Mice. Biochimica et Biophysica Acta (BBA) - Molecular and Cell Biology of Lipids 2014, 1841 (10), 1563–1570. https://doi.org/10.1016/j.bbalip.2014.08.003.

(38) Krings, A.; Rahman, S.; Huang, S.; Lu, Y.; Czernik, P. J.; Lecka-Czernik, B. Bone Marrow Fat Has Brown Adipose Tissue Characteristics, Which Are Attenuated with Aging and Diabetes. Bone 2012, 50 (2), 546–552. https://doi.org/10.1016/j.bone.2011.06.016.

(39) Schick, B. P.; Schick, P. K.; Chase, P. R. Lipid Composition of Guinea Pig Platelets and Megakaryocytes. The Megakaryocyte as a Probable Source of Platelet Lipids. Biochim Biophys Acta 1981, 663 (1), 239–248. https://doi.org/10.1016/0005-2760(81)90210-1.

(40) Long, M.; Williams, N. Immature Megakaryocytes in the Mouse: Morphology and Quantitation by Acetylcholinesterase Staining. Blood 1981, 58 (5), 1032–1039. https://doi.org/10.1182/blood.V58.5.1032.1032.

