## Supporting Information for "High Spatial Resolution MALDI Imaging Mass Spectrometry of Fresh-Frozen Bone"

CONTENTS

**Figure S1.** Autofluorescence image of a murine femur section without the use of Cryofilm……..……......**Page S-2**

**Figure S2.** Measurements of cracks in an autofluorescence image of a thaw-mounted femur section..….**Page S-3**

**Figure S3.** Lipid distributions by MALDI IMS at 10 µm spatial resolution with histological annotations.......**Page S-4**

**Figure S4.** Magnified images of megakaryocytes present in the bone marrow…………………...………....**Page S-5**

**Figure S5.** Enlarged ion images and replicate data for the matrix recrystallization experiment…………....**Page S-6**

**Table S1.** Additional MS parameters for the Bruker timsTOF flex……………………...……………………...**Page S-7**

**Table S2.** Additional MS parameters for the 15T Bruker solariX FT-ICR……………………...................**Page S-8**

**Table S3.** Mass accuracy of lipid identifications………………………………………………………………...**Page S-9**

**Table S4.** Mean intensity and increase values between glass and ITO slides..........................................**Page S-10**

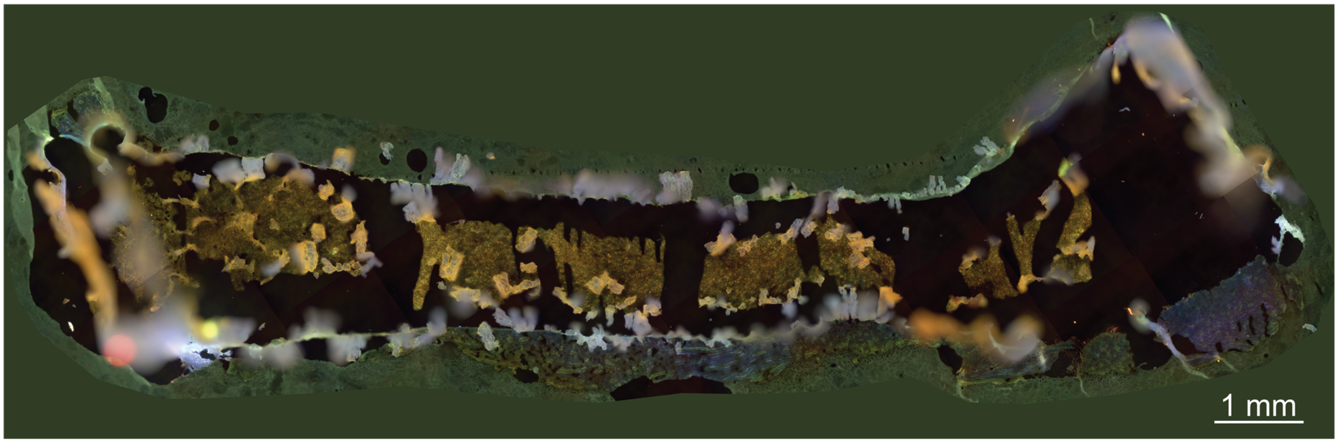


**Figure S1.** Cryosectioning a femur without Cryofilm leads to a notable collapse in tissue structure, as seen by the merged channel autofluorescence image.

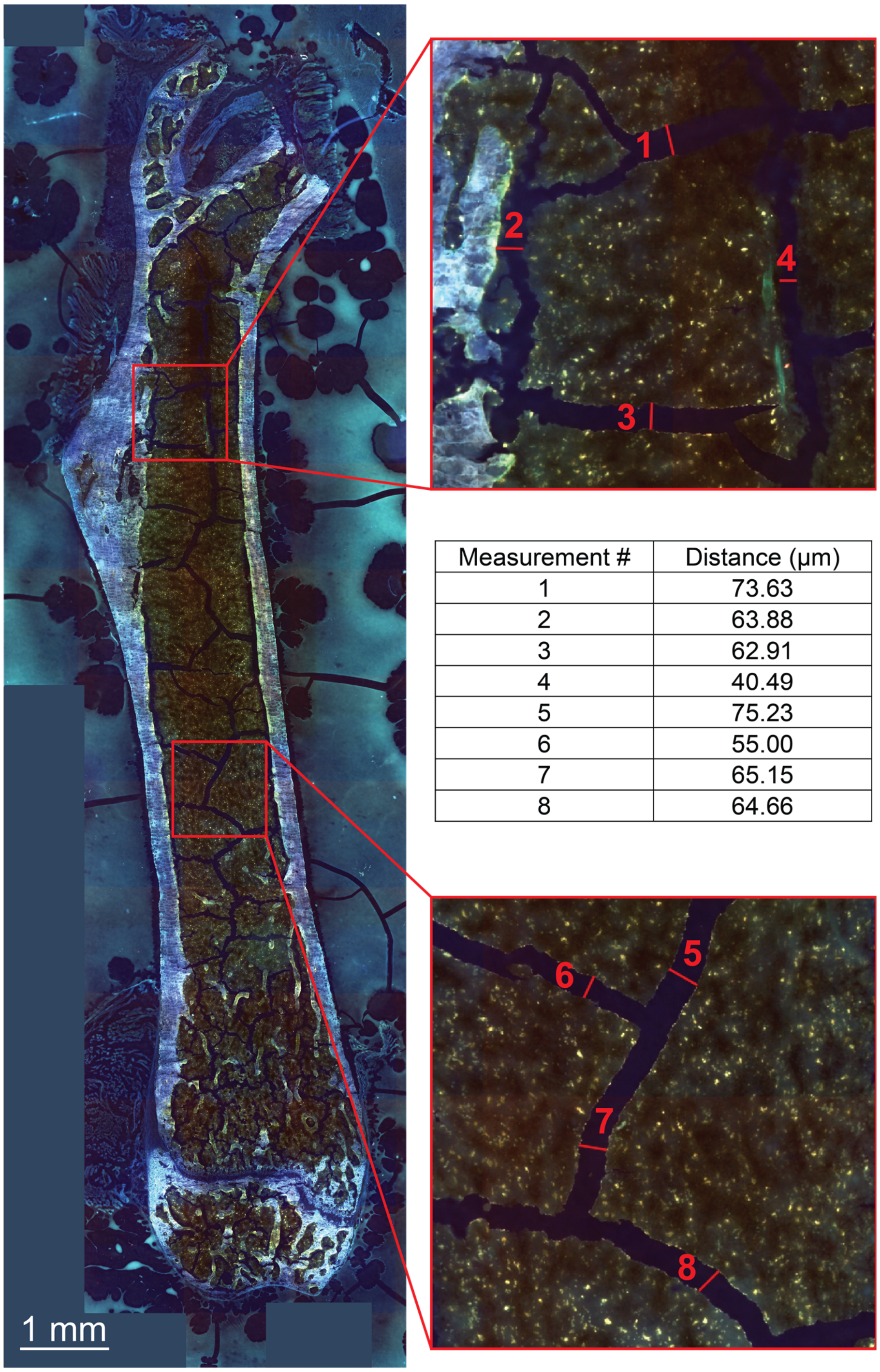


| Measurement # | Distance (µm) |
|---|---|
| 1 | 73.63 |
| 2 | 63.88 |
| 3 | 62.91 |
| 4 | 40.49 |
| 5 | 75.23 |
| 6 | 55.00 |
| 7 | 65.15 |
| 8 | 64.66 |

**Figure S2.** Thaw-mounting a femur section onto Cryofilm yields bone marrow cracks as large as 75 µm wide. Multiple measurements from an autofluorescence image are reported.

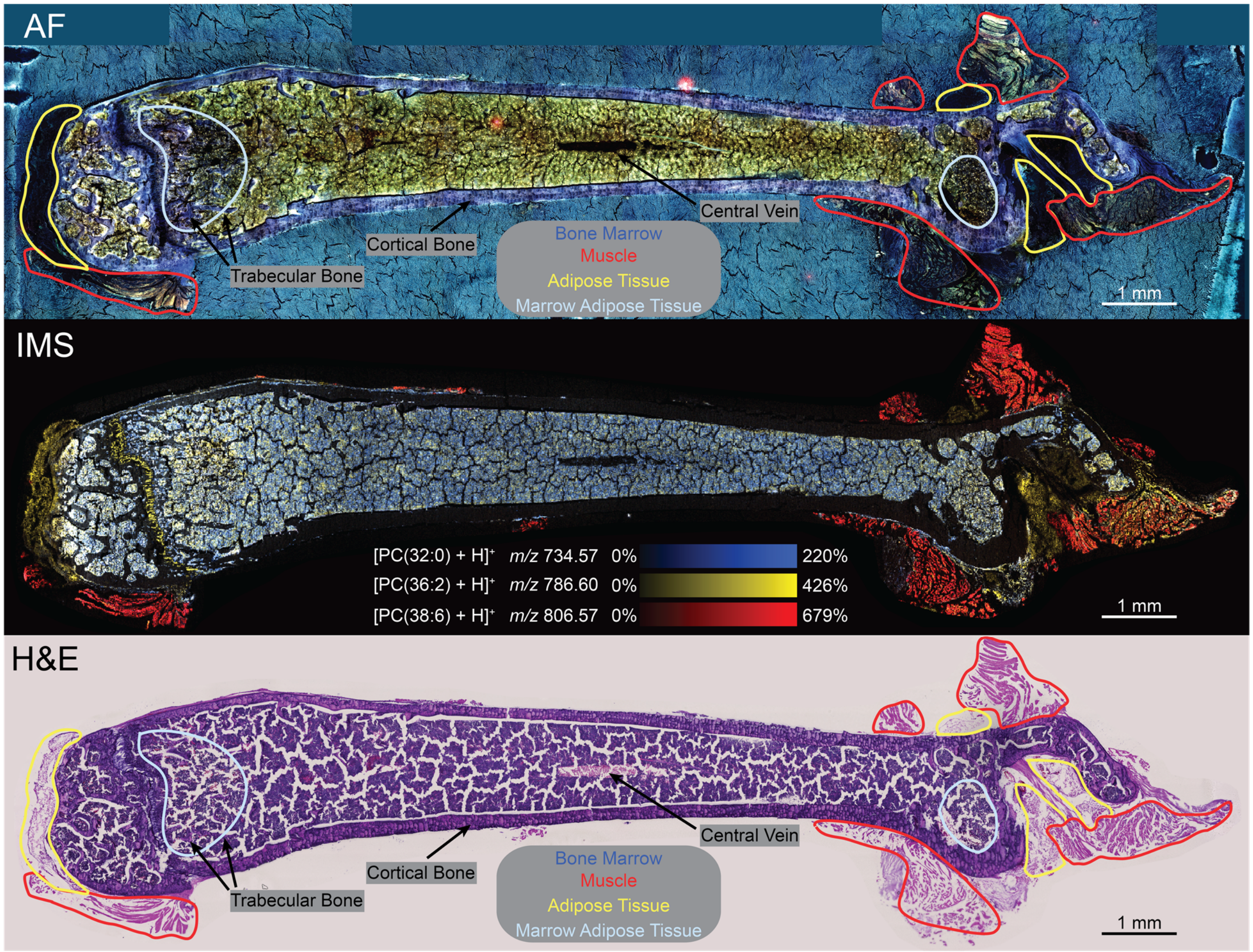


**Figure S3.** Sample preparation techniques enable imaging of a femur section at 10 µm spatial resolution. Bone marrow artifacts are accentuated in these ion images compared to 20 µm spatial resolution ion images. Similar to Figure 5, $[PC(32:0) + H]^+$ (*m/z* 734.57), $[PC(36:2) + H]^+$ (*m/z* 786.60), and $[PC(38:6) + H]^+$ (*m/z* 806.57) localize to bone marrow, adipose tissue, and muscle, respectively. Merged channel autofluorescence (pre-IMS) and H&E stain (post-IMS) images aid in histological annotations.

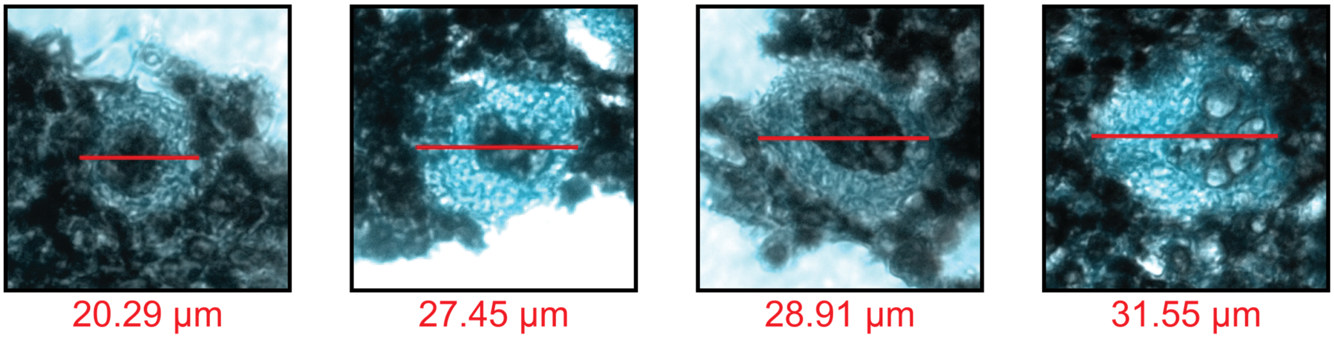


**Figure S4.** Megakaryocytes differ in cell diameter and nuclear configuration, which indicates variations in maturity. Images were acquired on an Axio Observer.Z1 microscope (Carl Zeiss Microscopy GmbH, Oberkochen, Germany) using a brightfield and DAPI channel.

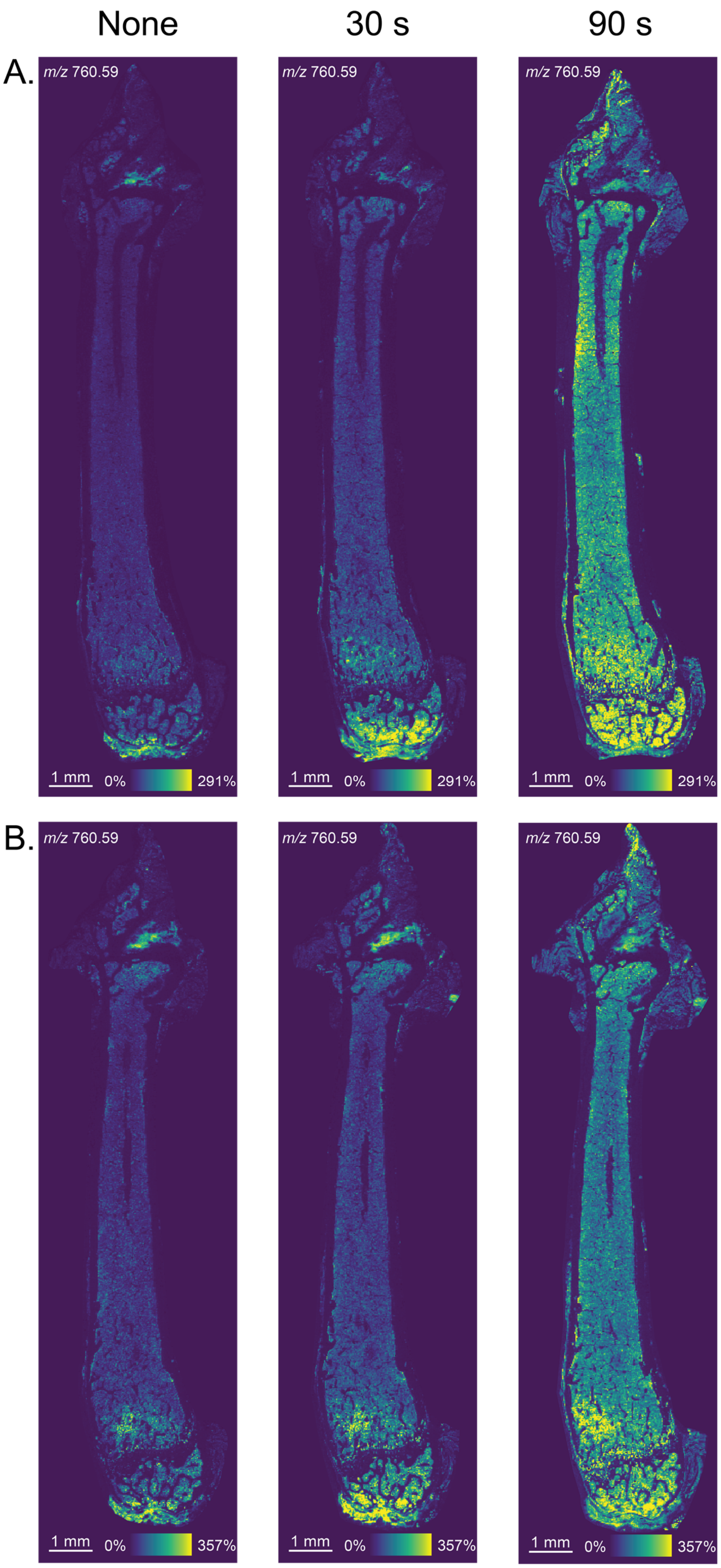


**Figure S5.** Ion images highlight a reproducible intensity increase that is dependent on the duration of matrix recrystallization. (A) Ion images from Figure 3 are enlarged. (B) Ion images are from a replicate experiment.

**Table S1.** Additional MS parameters are reported for the Bruker timsTOF flex.

| **Transfer** | |
|---|---|
| MALDI Plate Offset | 30.0 V |
| Deflection 1 Delta | 70.0 V |
| Funnel 1 RF | 450.0 Vpp |
| isCID Energy | 0.0 eV |
| Funnel 2 RF | 500.0 Vpp |
| Multipole RF | 500.0 Vpp |
| **Collision Cell** | |
| Collision Energy | 10.0 eV |
| Collison RF | 2900.0 Vpp |
| **Quadrupole** | |
| Ion Energy | 5.0 eV |
| Low Mass | 300.00 m/z |
| **Focus Pre TOF** | |
| Transfer Time | 110.0 µs |
| Pre Pulse Storage | 10.0 µs |

**Table S2.** Additional MS parameters are reported for the 15T Bruker solariX FT-ICR.

| **Source Optics** | | **Octopole** | |
|---|---|---|---|
| Capillary Exit | 220.0 V | Frequency | 5 MHz |
| Detector Plate | 200.0 V | RF Amplitude | 350.0 Vpp |
| Funnel 1 | 150.0 V | **Quadrupole** | |
| Skimmer 1 | 60.0 V | Q1 Mass | 610.00 m/z |
| Funnel RF Amplitude | 250.0 Vpp | **Collision Cell** | |
| **Transfer Optics** | | Collision Voltage | -1.0 V |
| Time of Flight | 0.850 ms | DC Extract Bias | 0.1 V |
| Frequency | 4 MHz | RF Frequency | 2 MHz |
| RF Amplitude | 380.0 Vpp | Collision RF Amplitude | 1200.0 Vpp |
| **Para Cell** | | **In Source Fragmentation** | |
| Transfer Exit Lens | -20.0 V | FSCID Collision Energy | 45.0 V |
| Analyzer Entrance | -10.0 V | **Shimming DC Bias** | |
| Side Kick | 2.0 V | 0°, 90°, 180°, 270° | 1.500 V |
| Side Kick Offset | -1.5 V | **Gated Injection DC Bias** | |
| Front Trap Plate | 1.500 V | 0°, 90°, 180°, 270° | 1.500 V |
| Back Trap Plate | 1.500 V | **MALDI Control** | |
| Back Trap Plate Quench | -30.0 V | Plate Offset | 100.0 V |
| Sweep Excitation Power | 18.0 % | Deflector Plate | 200.0 V |

**Table S3.** All lipids were identified with a <1 ppm mass error tolerance. Identifications for the most intense lipid species were made using LIPIDMAPS database (lipidmaps.org).

| ***m/z*** | ***m/z* (database)** | **ID (database)** | **Ion** | **Mass Error (ppm)** |
|---|---|---|---|---|
| 703.574285 | 703.5748 | SM 34:1; O2 | $[M+H]^+$ | 0.732 |
| 731.605766 | 731.6061 | SM 36:1; O2 | $[M+H]^+$ | 0.457 |
| 734.569636 | 734.5694 | PC 32:0 | $[M+H]^+$ | 0.321 |
| 760.584390 | 760.5851 | PC 34:1 | $[M+H]^+$ | 0.933 |
| 786.600759 | 786.6007 | PC 36:2 | $[M+H]^+$ | 0.075 |
| 787.668742 | 787.6687 | SM 40:1; O2 | $[M+H]^+$ | 0.053 |
| 806.568812 | 806.5694 | PC 38:6 | $[M+H]^+$ | 0.729 |
| 810.600437 | 810.6007 | PC 38:4 | $[M+H]^+$ | 0.324 |
| 813.684935 | 813.6844 | SM 42:2; O2 | $[M+H]^+$ | 0.658 |
| 815.699620 | 815.7000 | SM 42:1; O2 | $[M+H]^+$ | 0.466 |

**Table S4.** Cryofilm-bound tissue mounted to an ITO coated glass slide consistently improves the signal intensity of PCs and SMs. Mean intensity values (arbitrary) and intensity increase percentages are reported for multiple experiments. Replicate (Rep) #1 is represented in Figure 4. An ITO coating yields an intensity increase for all replicates, but the magnitude of improvement depends on the strength of the signal.

| Rep # - Slide | $[PC(32:0) + H]^+$ | $[PC(34:1) + H]^+$ | $[PC(36:2) + H]^+$ | $[PC(38:4) + H]^+$ | $[SM(34:1) + H]^+$ | $[SM(42:2) + H]^+$ |
|---|---|---|---|---|---|---|
| 1 - Glass | 2116.21 | 1731.74 | 645.59 | 325.577 | 451.695 | 110.221 |
| 1 - ITO | 3540.76 | 2938.13 | 1184.14 | 772.337 | 591.31 | 148.671 |
| **% Increase** | **67.32** | **69.66** | **83.42** | **137.22** | **30.91** | **34.88** |
| 2 - Glass | 2239 | 1551.72 | 506.618 | 246.92 | 222.719 | 103.957 |
| 2 - ITO | 4375.88 | 3315.24 | 1106.55 | 689.332 | 624.475 | 188.12 |
| **% Increase** | **95.44** | **113.65** | **118.42** | **179.17** | **180.39** | **80.96** |
| 3 - Glass | 539.924 | 101.741 | 33.3875 | 20.264 | 55.8127 | 6.7495 |
| 3 - ITO | 2375.72 | 386.127 | 160.549 | 132.174 | 389.533 | 50.8522 |
| **% Increase** | **340.01** | **279.52** | **380.87** | **552.26** | **597.93** | **653.42** |
